# Reliability of quantitative multiparameter maps is high for MT and PD but attenuated for R1 and R2* in healthy young adults

**DOI:** 10.1101/2021.11.10.467254

**Authors:** Elisabeth Wenger, Sarah E. Polk, Maike M. Kleemeyer, Nikolaus Weiskopf, Nils C. Bodammer, Ulman Lindenberger, Andreas M. Brandmaier

## Abstract

We investigate the reliability of individual differences of four quantities measured by magnetic resonance imaging based multiparameter mapping (MPM): magnetization transfer (MT), proton density (PD), longitudinal relaxation rate (R1), and effective transverse relaxation rate (R2*). A total of four MPM datasets, two on each of two consecutive days, were acquired in healthy young adults. On Day 1, no repositioning occurred; on Day 2, participants were repositioned between MPM datasets. Using intra-class correlation effect decomposition (ICED), we assessed the contributions of session-specific, day-specific, and residual sources of measurement error. For whole-brain gray and white matter, all four MPM parameters showed high reproducibility and high reliability, as indexed by the coefficient of variation (CoV) and the intra-class correlation (ICC). However, MT, PD, R1, and R2* differed markedly in the extent to which reliability varied across brain regions. MT and PD showed high reliability in almost all regions. In contrast, R1 and R2* showed low reliability in some regions outside the basal ganglia, such that the sum of the measurement error estimates in our structural equation model was higher than estimates of between-person differences. In sum, in this sample of healthy young adults, the four MPM parameters showed very little variability over four measurements over two days but differed in how well they could assess between-person differences. We conclude that R1 and R2* might carry only limited person-specific information in samples of healthy young adults, and, by implication, might be of restricted utility for studying associations to between-person differences in behavior.

## Introduction

Research on human development seeks to delineate the variable and invariant properties of age-graded changes in the organization of brain-behavior-environment systems (Lindenberger, Li, & Bäckman, 2006). Magnetic resonance imaging (MRI) has become an indispensable tool for the noninvasive assessment of brain anatomy and microstructure and will continue to contribute knowledge on how brain structure changes in response to new environmental challenges or aging.

Quantitative MRI can help us to characterize the brain’s microanatomy by using the magnetophysical properties of water molecules in brain tissue that govern MRI contrasts, which are then in turn used as surrogate parameters to describe histological properties (Tofts, 2003; Weiskopf, Edwards, Helms, Mohammadi, & Kirilina, 2021). Recently, a comprehensive quantitative multiparameter mapping approach was developed, which provides high resolution maps of the longitudinal relaxation rate (R_1_ = 1/T_1_), proton density (PD), magnetization transfer saturation (MT) and effective transverse relaxation rate (R_2_* = 1/T_2_*) (Helms, Dathe, & Dechent, 2008; Helms, Draganski, Frackowiak, Ashburner, & Weiskopf, 2009; Weiskopf et al., 2011). These multiparameter maps are related to microstructural properties of myelin, iron deposits, and water, amongst other things (Draganski et al., 2011), even though it is not a simple one-to-one mapping and the exact relation to underlying physiological processes at the cellular and molecular level is still to be resolved (Weiskopf et al., 2021).

Central questions in lifespan psychology often pertain to the range and direction of within-person change and variability, be it longitudinal change observed over years and decades (Raz & Rodrigue, 2006), intervention-induced change over weeks and months (May, 2011), or fluctuations that occur from day to day and from moment to moment (Schmiedek, Lövdén, & Lindenberger, 2010). Random measurement error and systematic drifts can compromise the reliable measurement of change (Karch et al., 2019). Given that the expected effect sizes we typically seek to detect with structural MR are often no larger than 2 to 3% of the quantity under investigation, be it gray matter volume, mean diffusivity, or other structural brain measures, measurement error and drift can easily jeopardize the reliable assessment of within-person changes and between-person differences. If measurement artefacts are of similar magnitude as effects of interest, then reliability is low, and effects of interest cannot be detected. Reliability is a pivotal issue in longitudinal studies, but it also matters for crosssectional studies, when researchers either are interested in stable between-person differences or when they use time- or age-related differences between people as a proxy for change. Thus, in both cross-sectional and longitudinal designs, the stability of MR measures cannot simply be assumed, but must instead be tested explicitly (Noble, Scheinost, & Constable, 2020).

Different scientific communities such as physics and psychometrics can rely on two fundamentally different conceptions of reliability and error: coefficient of variation (CoV) and intra-class correlation coefficient (ICC). Each of them is equally important but notably provide answers to very different questions. Physicists typically inquire how reliably a given measurement can detect a given quantity. Therefore, it is common and well-justified to use CoV as the main measure to assess repeatability, as is done for example in a previous multicenter study of MPM (Weiskopf et al., 2013) or another study testing within-site and between-site reproducibility (Leutritz et al., 2020) of MPM. The CoV is a standardized measure of dispersion and is often expressed as a percentage. It is widely used in analytical chemistry, engineering, and physics to express precision of a measurement and repeatability on well-defined objects of measurement. However, Brandmaier, Wenger, and colleagues (2018) showed that CoV does not distinguish between error variance and true construct-related variance, that is, between-person differences in the construct of interest, and may therefore not be particularly informative for correlational studies interested in assessing and explaining between-person differences. Instead, cognitive neuroscience commonly relies on a different conception of reliability, which refers to the precision of assessing between-person differences. This is typically expressed as a ratio index, the ICC, which relates the variance within persons (or groups of persons) to the total variance, and therefore represents the strength of association between any pair of measurements made on the same object (Bartko, 1966). It is important to keep in mind that ICC will increase when within-subject measurements become more similar or when the true scores of participants become more distinct from one another. By definition, ICC values must thus be interpreted contingent upon the characteristics of a given population.

To assess the adequacy of MPM parameters for correlational studies of human neuroscience, we investigated the reliability of MPM parameters within participants across four different measurement occasions in relation to between person-differences, using ICC. Specifically, we made use of intra-class correlation effect decomposition (ICED), which has been recently introduced by (Brandmaier, Wenger, et al., 2018). ICED estimates overall reliability while attributing the overall error variance to different sources by making use of the design features of a given study. We acquired data from 15 volunteers, who each were assessed four times. On Day 1, participants were measured twice back-to-back, without repositioning between MPM datasets. On Day 2, participants were also measured twice, but this time with a break in between measurements, which afforded repositioning of the participants’ head. With this study design, we are able to tease apart three sources of error variance: variance originating from (1) repositioning the subject between two measurements on the same day (session-specific error variance; mostly due to different head position inside the coil); (2) repositioning the subject on another day (day-specific error variance; e.g., due to different environmental properties, intra-subject changes, or scanner-related properties); (3) other sources of error (residual error variance). Given that the quantities derived from multiparameter maps, if measured reliably, can contribute to a better understanding of individual differences in brain physiology and age-related changes therein, estimating the size of these sources of error variance is of great methodological interest. In particular, if the sum of these three sources of error is small relative to the magnitude of between-person differences, then reliability as indexed by ICC is high, which bodes well for the investigation of individual differences in brain physiology and potential relations to individual differences in behavior. Conversely, is if the sum of these three sources of error is relatively large, then these parameters are not well suited for investigating individual differences of any sort, including associations to behavior.

## Methods

### Participants and Procedure

Fifteen healthy volunteers (8 females, mean age = 27.30, SD = +/- 3.34, range = 22 – 31 years) participated in the study. All participants had normal hearing, normal or corrected-to-normal vision, no history of psychological or neurological diseases, and no contraindication to participate in an MR study, like metallic implants, tinnitus, or claustrophobia. The sample was quite representative regarding general cognitive functioning, as indicated by perceptual speed performance measured via the Digit Symbol substitution test (Wechsler, 1981; M = 64.5, SD = 6.9) and vocabulary via the Spot-a word test (MWT-A, Lehrl, Merz, Burkard, & Fischer, 1991; M = 31.3, SD = 2.5) (see Figure 1). For comparison, data from a meta-analysis by Hoyer and colleagues (2004) showed a mean performance in the Digit Symbol substitution test of 69.8 in younger adults, and data from another Berlin based training study with 100 younger adults had a mean performance of 60.3 (SD = 9.5) in this test (Schmiedek et al., 2010), and data from yet another Berlin based training study with 44 younger adults reported a very comparable mean performance in the vocabulary test of 30.3 (SD = 2.7) (Lövdén et al., 2012). These behavioral test results give reason to believe that the chosen sample is representative of a young adult population on the cognitive level, and may therefore most likely also show to-be-expected variance in brain structure, even though this link from cognition to brain structure is of course speculative.

**Figure 1.**
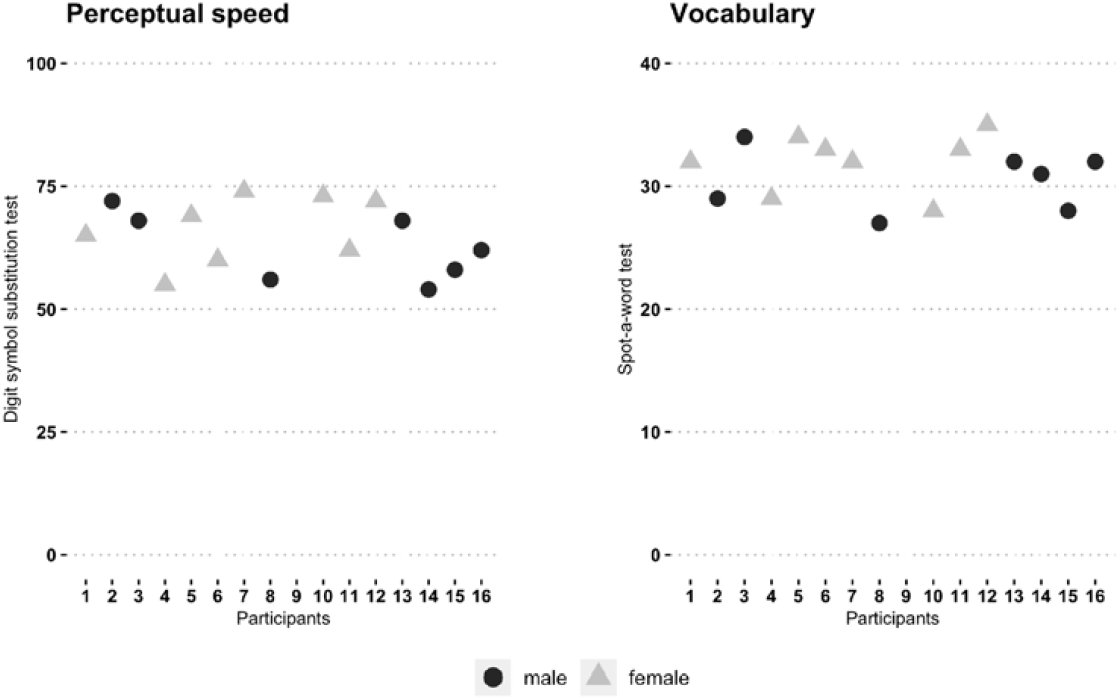
General cognitive functioning of all 15 participants, as measured by the Digit symbol substitution test for perceptual speed (Wechsler, 1981; M = 64.5, SD = 6.9) and the Spot-a-word test (MWT-A) for vocabulary knowledge (Lehrl, Merz, Burkard, & Fischer, 1991; M = 31.3, SD = 2.5).

Each participant was scanned four times, distributed over two consecutive days. On Day 1, participants were scanned for the first time with the full MPM protocol (Measurement 1). No repositioning was done, i.e., participants remained inside the scanner, but all scanner adjustments and settings were reset. Then, participants were measured a second time with the MPM protocol (Measurement 2). For the Day 2 measurement, all participants were re-invited to be scanned on the following day around the same time of the day and the full MPM protocol was acquired again (Measurement 3). After that, participants were moved out of the scanner, the head coil was removed and participants briefly got up and walked around before lying back down on the scanner bed and being moved back in. Participants were then scanned for the fourth time (Measurement 4).

The study received ethical approval by the ethics committee of the German Association of Psychology (Deutsche Gesellschaft für Psychologie, DGPs) and was carried out as a pilot study in preparation for a training intervention study. All participants provided written informed consent prior to participation.

### MR Image Acquisition

All MPM datasets were acquired on a Siemens Tim Trio 3T MR scanner (Erlangen, Germany; VB17a software version) with a standard radio-frequency (RF) 32-channel receive head coil and RF transmit body coil. The MPM protocol comprised one static magnetic (B_0_) GRE-field map, one RF transmit field map (B_1_^+^), and three multi-echo 3D FLASH (fast low angle shot) sequences. The MPM acquisition and post-processing have been developed and described in previous studies (Helms, Dathe, & Dechent, 2008; Helms, Dathe, Kallenberg, & Dechent, 2008; Weiskopf et al., 2011, 2013) and acquisition parameters were chosen in accordance with previously published work (Weiskopf et al., 2013). Thus, they are only described in brief here.

The B_0_ gradient echo field mapping sequence was acquired with the following parameters: 64 transverse slices, slice thickness = 2 mm with 50% distance factor, repetition time (TR) = 1020 ms, echo times (TE) TE1/TE2 = 10/12.46 ms, flip angle α = 90°, matrix = 64 × 64, field of view (FOV) = 192 × 192 mm, right-left phase encoding (PE) direction, bandwidth (BW) = 260 Hz/Px, flow compensation, acquisition time = 2:14 min.

Maps of the local RF transmit/B_1_^+^ field were acquired following recommendations by Lutti and colleagues (Lutti, Hutton, Finsterbusch, Helms, & Weiskopf, 2010) and were measured and estimated from a 3D EPI acquisition of spin and stimulated echoes (SE and STE) with different flip angles. The following parameters were used: 4 mm isotropic resolution, matrix = 64 × 48 × 48, FOV = 256 × 192 × 192 mm, parallel imaging using GRAPPA factor 2 × 2 in PE and partition directions, TR = 500 ms, TE_SE_/_STE_/mixing time = 39.06 ms/33.80 ms. Eleven pairs of SE/STE image volumes were measured successively employing decreasing flip angles α from 115° to 65° in steps of −5° (applied in a α–2α–α series of RF pulses to produce SEs and STEs; see Akoka, Franconi, Seguin, & Le Pape, 1993). Acquisition time was 3 min.

The three different multi-echo FLASH sequences were acquired with predominantly T_1_ weighting (T_1_w), proton density weighting (PDw), or magnetization transfer weighting (MTw) by appropriate choice of repetition time (TR) and flip angle α (T_1_w: TR/α = 24.5 ms/21°; PDw and MTw: TR/α = 24.5 ms/6°) and by applying an off-resonance Gaussian-shaped RF pulse (4 ms duration, 220° nominal flip angle, 2 kHz frequency offset from water resonance) prior to excitation in case of the MTw sequence version. Multiple gradient echoes with alternating readout polarity were acquired at six equidistant echo times (TE) between 2.34 ms and 14.04 ms for the T_1_w and MTw acquisitions with two additional echoes at TE = 16.38 ms and 18.72 ms for the PDw acquisition. A high readout bandwidth BW = 465 Hz/pixel was used to minimize off-resonance artifacts. To speed up data acquisition, GRAPPA parallel imaging with an acceleration factor of two was applied in the phase-encoding (anterior-posterior) direction (outer/slow phase encoding loop) and 6/8 partial Fourier acquisitions in the partition (left-right) direction (inner/fast phase encoding loop). Additional acquisition parameters were as follows: 1 mm isotropic resolution, 176 slices per slab, FOV = 256 × 240 mm, acquisition time of each of the three FLASH sequences = 7:03 min.

### Estimation of Parameter Maps

All data analyses and processing were performed in SPM12 (www.fil.ion.ucl.ac.uk/spm) running on Matlab 2017b (The MathWorks Inc., Natick, MA, USA) using the hMRI toolbox (Tabelow et al., 2019; https://hmri-group.github.io/hMRI-toolbox/). The *Create hMRI maps* module was used to compute quantitative as well as semi-quantitative estimates of R2*, R1, PD, and MT from unprocessed multi-echo T1-, PD-, and MT-weighted RF-spoiled gradient echo acquisitions.

As has been described in more detail elsewhere (Helms, Dathe, & Dechent, 2008; Helms, Dathe, Kallenberg, et al., 2008; Weiskopf et al., 2011, 2013), the signal from the PD-, and T_1_-weighted echoes can be described by the Ernst equation, whereas the signal strength throughout a train of gradient-recalled echoes follows a largely exponential decay with time constant T_2_* – for all three contrasts. For the MT-weighted contrast Helms et al. have suggested to treat the MT-weighting preparation of the sequence like a first pulse in a dual-excitation FLASH sequence, and introduced – based on the associated extended Ernst equation – a novel semi-quantitative parameter for describing the MT saturation effect (Helms, Dathe, & Dechent, 2008). This novel MT parameter is ‘semi-quantitative’ since it still depends on the efficiency of the applied MT saturation, but – different from the frequently-used magnetization transfer ratio (MTR) – an influence by the local T_1_ and also by transmit field inhomogeneities is largely cancelled.

The R2*, i.e., the effective transverse relaxation rate (R2* = 1/T_2_*) was estimated by applying the ESTATICS approach (Weiskopf, Callaghan, Josephs, Lutti, & Mohammadi, 2014) assuming mono-exponential signal decay with increasing TE with the same R2* for all three contrast weightings. That is, for all three contrasts (PDw, T1w and MTw) – and within each contrast for the number of available TEs as datapoints – a joint log-linear fit using ordinary least squares (OLS) is applied. Thereby the slope corresponds to R2*, identically for all three contrasts, whereas the intercept, i.e. the signal values extrapolated to TE = 0, are representing the three different contrasts without any influence of this common transversal relaxation. With this approach relatively stable values for R2* are estimated for each voxel; additionally, and values largely unaffected by R2* are estimated for all three contrasts (i.e. extrapolated to TE=0). Due to their minimized dependency on R2*, these images are an optimal basis for further calculations.

As a next step, uncorrected R1, PD, and MT maps are calculated from the extrapolated T1w-, PDw, and MTw measurements (for TE = 0) by applying the Ernst equation according to Helms and colleagues (Helms, Dathe, & Dechent, 2008; Helms, Dathe, Kallenberg, et al., 2008). Quantitative maps of R1, i.e., the longitudinal relaxation rate (R1 = 1/T_1_) were corrected for local RF transmit field inhomogeneities. To do so, the acquired 3D-EPI-based B1^+^ maps (Lutti et al., 2012) were used after correcting them for EPI-specific distortions by means of the gradient echo-based B0 fieldmaps. Also the MT maps were corrected for residual local RF transmit field inhomogeneities – using a semi-empirical approach (Rowley et al., 2021; Weiskopf et al., 2013). Imperfect RF spoiling was also corrected for in the T_1_ maps (Preibisch & Deichmann, 2009). PD maps were estimated from the signal amplitude maps by adjusting for global and local receive sensitivity inhomogeneities using the “unified segmentation” approach (Ashburner & Friston, 2005). The mean white matter PD value was calibrated to 69 percent units, since the global mean PD cannot be estimated accurately without an external standard.

To achieve an improved within-subject coregistration of the created maps, we adapted a longitudinal processing pipeline of the data. To do this, we first thresholded all MT maps (between 0 and 5) and all PD maps (between 0 and 200) in order to improve segmentation performance. We then conducted multichannel segmentations using the thresholded MT and PD map pairs for each measurement and each subject. The resulting gray and white matter segments from the four measurements per subject – formatted to get imported to DARTEL (option ‘DARTEL imported’) – were the input to SHOOT to create an unbiased within-subject registration. The respective deformations were applied to the raw MT and PD maps to warp them into each subject’s template space, and the median MT and PD map across all measurements of one individual was computed. These median maps were (again) subjected to a multichannel segmentation and a group template was created from the resulting ‘DARTEL imported’ gray and white matter segments of all subjects using again SHOOT. Subsequently, we combined the two deformation fields, namely the one from native space to the subject’s template space and the second from the subject’s template space to group template space. For each measurement, this combined deformation field together with all four parameter maps (MT, PD, R1, R2*) was fed into SHOOT normalize to achieve normalization to MNI space. Also, the gray and white matter segments derived by the multichannel segmentation of the median MT and PD maps were spatially normalized to MNI space (Jacobian modulation was applied in this case). From these normalized tissue class segments together with the four parameter maps finally ‘*smoothed tissue specific MPMs*’ were computed applying a 6mm FWHM smoothing kernel and weighted averaging (the full set of scripts can be found on https://git.mpib-berlin.mpg.de/plasticity/aktiv/hmri_scripts.git).

These segmented modulated maps were used to extract the mean and standard deviation values across voxels in each one of the pre-defined regions of interest (ROIs) for every individual, at every measurement time point from the Harvard-Oxford cortical and subcortical structural atlases (https://identifiers.org/neurovault.collection:262; Desikan et al., 2006). We focused on a selection of ROIs that are typically of interest to neuroscientists when for example investigating language learning and effects of physical exercise. We therefore extracted means and SD for whole gray matter (cortex; GM), whole white matter (WM), inferior frontal gyrus (IFG) pars triangularis (pars tri), IFG pars opercularis (pars oper), orbitofrontal cortex (OFC), anterior cingulate cortex (ACC), precuneus, middle temporal gyrus (MTG), caudate nucleus, putamen, and pallidum (see Figure 2). Additionally, we also performed a whole-brain voxel-wise analysis, such that reliability was also estimated for every individual voxel.

**Figure 2.**
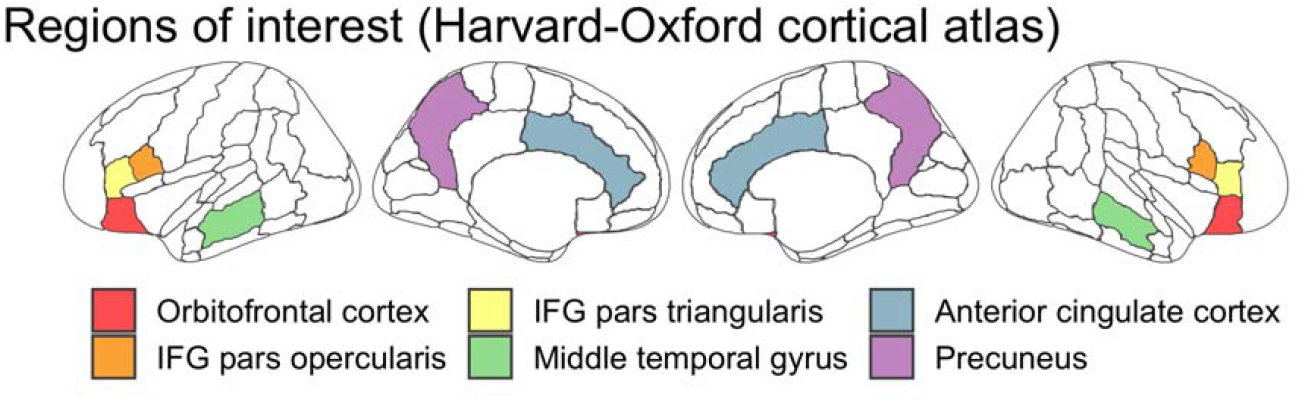
Cortical regions of interest (ROIs) from the Harvard-Oxford atlas. In addition to the six selected regions displayed here, we also investigated the subcortical regions caudate, putamen, and pallidum, as well as whole gray matter and white matter Additionally, we investigated reliability in a voxel-wise manner.

### Statistical Analysis

We used intra-class effect decomposition (ICED) to estimate reliability (Brandmaier, Wenger, et al., 2018). This recently introduced approach uses structural equation modeling (SEM) of data to decompose reliability in orthogonal sources of measurement error that can be attributed to different measurement characteristics. Using ICED, we are able to estimate the main effects of session, day, and residual variance on measurement error. It therefore allows to distinguish between the following variance compartments: true-score variance (Var T; representing true between-person differences in the construct of interest), day-specific error variance (Var D), session-specific error variance (Var S; here capturing the effect of repositioning a person between the MPM datasets, accompanied by a new prescan, i.e. new adjustments of RF amplitude, center frequency and B0 shim), and residual error variance (Var E). Model specification and estimation were performed in Ωnyx (von Oertzen, Brandmaier, & Tsang, 2015) and lavaan, an SEM package for the statistical programming language R (Rosseel, 2012).

The path diagram in Figure 3 illustrates the ICED model for estimating the individual variance components of the total observed variance of the MPM parameter magnetization transfer saturation (MT) in gray matter. The four measurements are labeled in the path diagram as “GM_MT_D1M1”, “GM_MT_D1M2”, “GM_MT_D2M1”, and “GM_MT_D2M2”. The same labeling convention was applied to all parameters (MT, PD, R1, R2*) in all ROIs and in every voxel.

**Figure 3.**
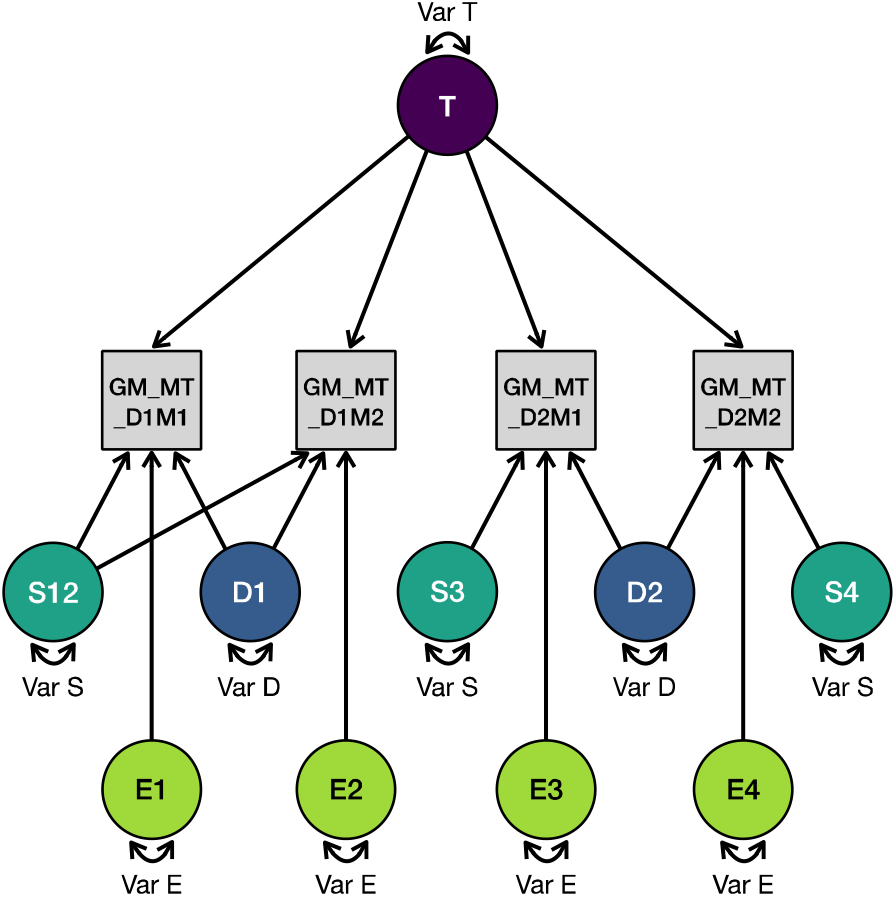
Path diagram of a structural equation model derived from Ωnyx. This diagram exemplifies the parameter magnetization transfer saturation (MTsat) in whole gray matter (GM). In our repeated measures design, each participant was scanned four times: twice on Day 1 without repositioning (D1M1 & D1M2), and twice on Day 2 with repositioning (D2M1 & D2M2). Data were standardized across days and measurements, and a saturated mean structure was used. Var T = true score variance, Var S = session-specific variance, Var D = day-specific variance, Var E = residual error variance.

**Figure 4.**
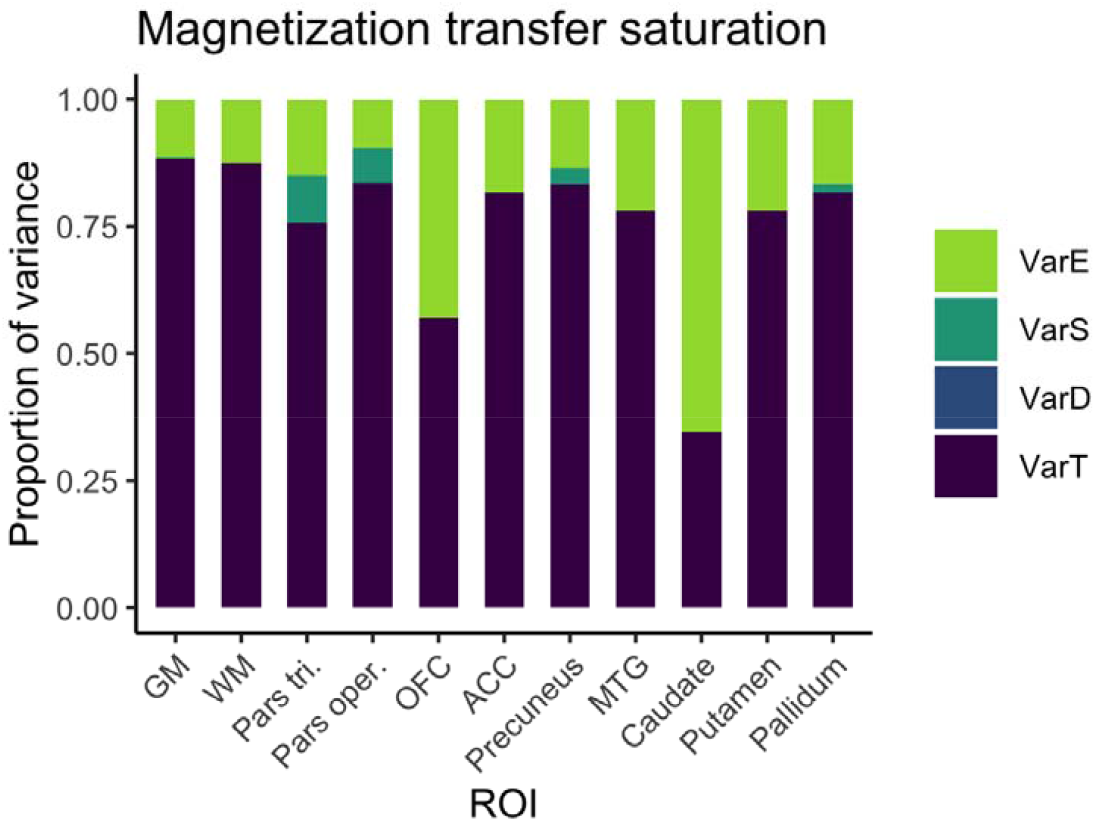
Distribution of magnitudes of sources of variance for magnetization transfer across the different ROIs. Note that absolute magnitudes are displayed, irrespective of significance. Session- and day-specific variances were not significantly different from zero for MT.

**Figure 5.**
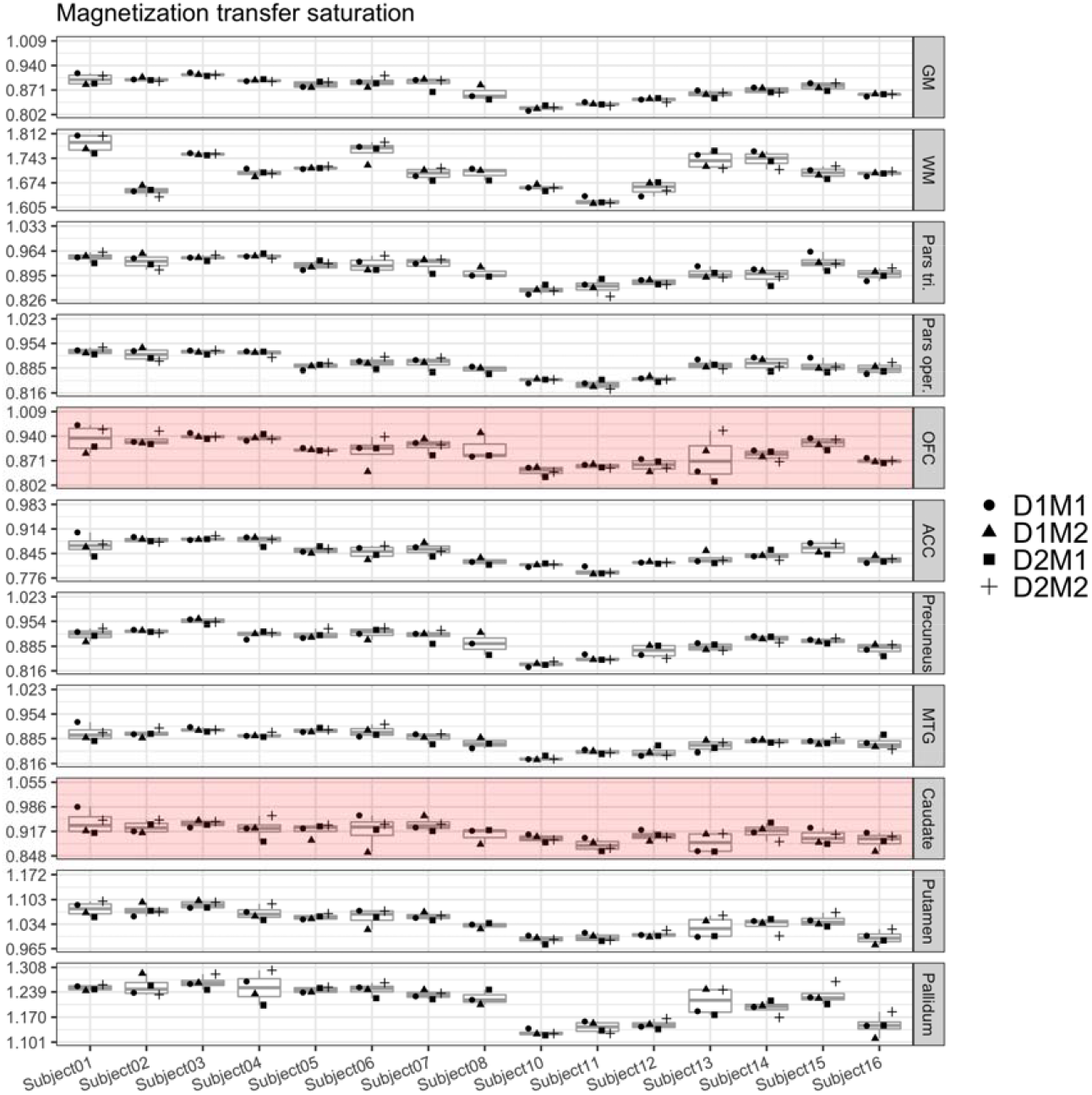
Boxplots of MT values for each of the 15 participants and their four measurements. Parts of the plots marked in red indicate an attenuated ICC value (<0.75) for this region. Note that the range displayed on the y-axis differs across ROIs; importantly, though, the width of the displayed range is constant across ROIs to ensure comparability and is always 0.207.

**Figure 6.**
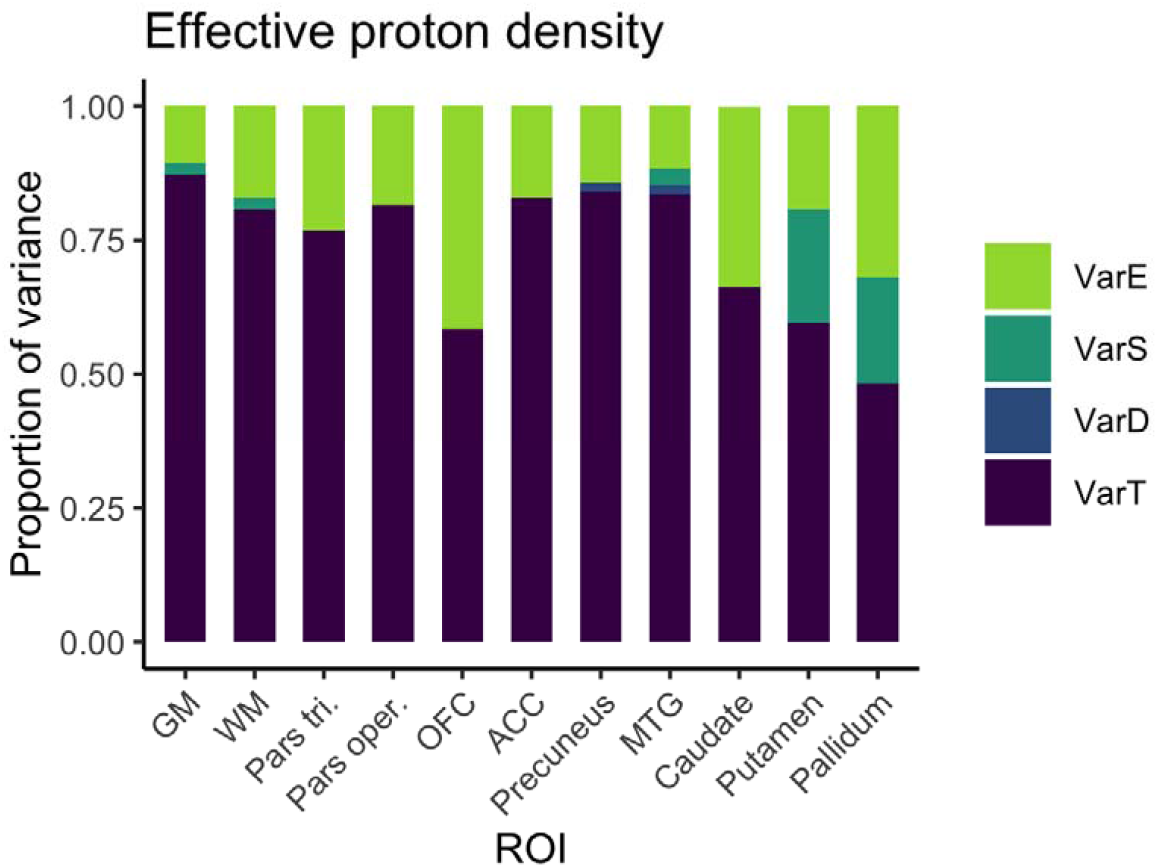
Distribution of magnitudes of sources of variance for PD across the different ROIs. Note that absolute magnitudes are displayed, irrespective of significance. Session- and day-specific variances were not significantly different from zero for PD.

First, the baseline SEM model was generated, which estimated all four variance parameters related to the four sources of variance (Var T, Var D, Var S, Var E), with a lower bound of 0.0001 applied to the estimates. Three additional null models were generated in which the true, day-related, and session-related variance were respectively set to zero, one at a time. To assess the significance of the magnitudes of these separate sources of error, likelihood ratio tests were used to compare the unconstrained models against the respective null models. A Wald test was used to test the residual error variance component, as the null model without an orthogonal error structure cannot be estimated. We report all variances rescaled such that they add up to one; this way, the variances can directly be interpreted as relative contributions to overall variance or variance explained.

These variance components were then used to calculate ICC, an index of reliability between MPM datasets within the study. We defined ICC as the ratio of between-person variance to total variance at the level of observed variables. In addition, we calculated ICC2 as a measure of reliability on the construct level. As such, it is defined as the ratio of true score variance to total (effective) variance, where the effective error is the single residual error term that arises from all variance components other than the construct that is to be measured. When assuming no day-specific and session-specific effects, we would obtain exactly the classical definition of ICC2, which scales the residual error variance with the number of measurement occasions (please see Brandmaier et al., 2018, for exact definitions and formulae). In sum, ICC is a coefficient describing test-retest reliability of a single measurement (i.e., how well can a single measurement measure the underlying quantitative value), whereas ICC2 is a coefficient describing test-retest reliability of the entire design (i.e., how well can we measure the underlying quantitative value with multiple measurements; here a total of 4 measurements in the given study design. As ICC2 is contingent upon a certain study design (with a specific number of measurement occasions), we prefer to rely on ICC in our interpretation of the data, as it relies on a single occasion of measurement per person and thus provides a lower bound of reliability. ICC2 values are additionally reported for reference. Note that as the variance estimates were rescaled such that they represent proportions of the total variance, true score variance is equal to the ICC here.

Bootstrapped 95% confidence intervals, using 1000 samples, were generated for ICC and ICC2 values of each parameter (MT, PD, R1, R2*) in all ROIs (using *boot.ci* in lavaan).

We report mean values of all MPM parameters as well as CoVs. We calculated CoV for each parameter in each ROI by dividing the standard deviation of the four extracted means (SD, normalized by N–1 sample size to avoid bias; in our case 4 scans – 1) by the overall mean across all 4 measurement points (CoV = SD/Mean). In addition, we visualize true score variance, that is ICC, in every voxel in whole-brain maps where the whiter a voxel is displayed, the closer its ICC is to 1.

## Results

### Mean MPM parameter values and CoVs

The means and SDs, as well as CoVs of all four MPM parameters in all ROIs are presented in Table 1 and 2.

**Table 1.** Means and standard deviation for each ROI and for each MPM parameter: magnetization transfer saturation (MTsat), proton density (PD), longitudinal relaxation rate (R1), and effective transverse relaxation rate (R2*).

| ROIs | MTsat (p.u.) | PD (p.u.) | R1 ($\text{s}^{-1}$) | R2* ($\text{s}^{-1}$) |
| --- | --- | --- | --- | --- |
| Gray matter | $0.874 \pm 0.011$ | $80.61 \pm 0.26$ | $0.622 \pm 0.007$ | $16.9 \pm 0.2$ |
| White matter | $1.706 \pm 0.017$ | $69.69 \pm 0.10$ | $0.967 \pm 0.012$ | $21.5 \pm 0.2$ |
| IFG (pars tri.) | $0.912 \pm 0.016$ | $80.11 \pm 0.38$ | $0.644 \pm 0.011$ | $16.2 \pm 0.4$ |
| IFG (pars oper.) | $0.893 \pm 0.013$ | $80.46 \pm 0.33$ | $0.629 \pm 0.010$ | $15.9 \pm 0.4$ |
| OFC | $0.898 \pm 0.027$ | $79.96 \pm 0.58$ | $0.635 \pm 0.016$ | $17.5 \pm 0.4$ |
| ACC | $0.846 \pm 0.014$ | $81.98 \pm 0.29$ | $0.598 \pm 0.012$ | $14.9 \pm 0.4$ |
| Precuneus | $0.899 \pm 0.014$ | $80.57 \pm 0.27$ | $0.615 \pm 0.012$ | $17.5 \pm 0.5$ |
| MTG (posterior) | $0.879 \pm 0.013$ | $80.53 \pm 0.37$ | $0.617 \pm 0.008$ | $16.7 \pm 0.3$ |
| Caudate | $0.913 \pm 0.024$ | $81.31 \pm 0.40$ | $0.684 \pm 0.019$ | $19.2 \pm 0.6$ |
| Putamen | $1.039 \pm 0.018$ | $79.26 \pm 0.44$ | $0.744 \pm 0.013$ | $21.3 \pm 0.5$ |
| Pallidum | $1.211 \pm 0.023$ | $77.39 \pm 0.59$ | $0.842 \pm 0.017$ | $29.7 \pm 0.7$ |

*Note.* Values extracted for ROIs from the Harvard-Oxford atlas, from the longitudinally processed normalized and segmented parameter maps. Means were calculated across all voxels and all participants. Standard deviations were calculated across the means of the four measurement points (normalized by $n-1=3$ ). p.u. = percentage units.

**Table 2.**
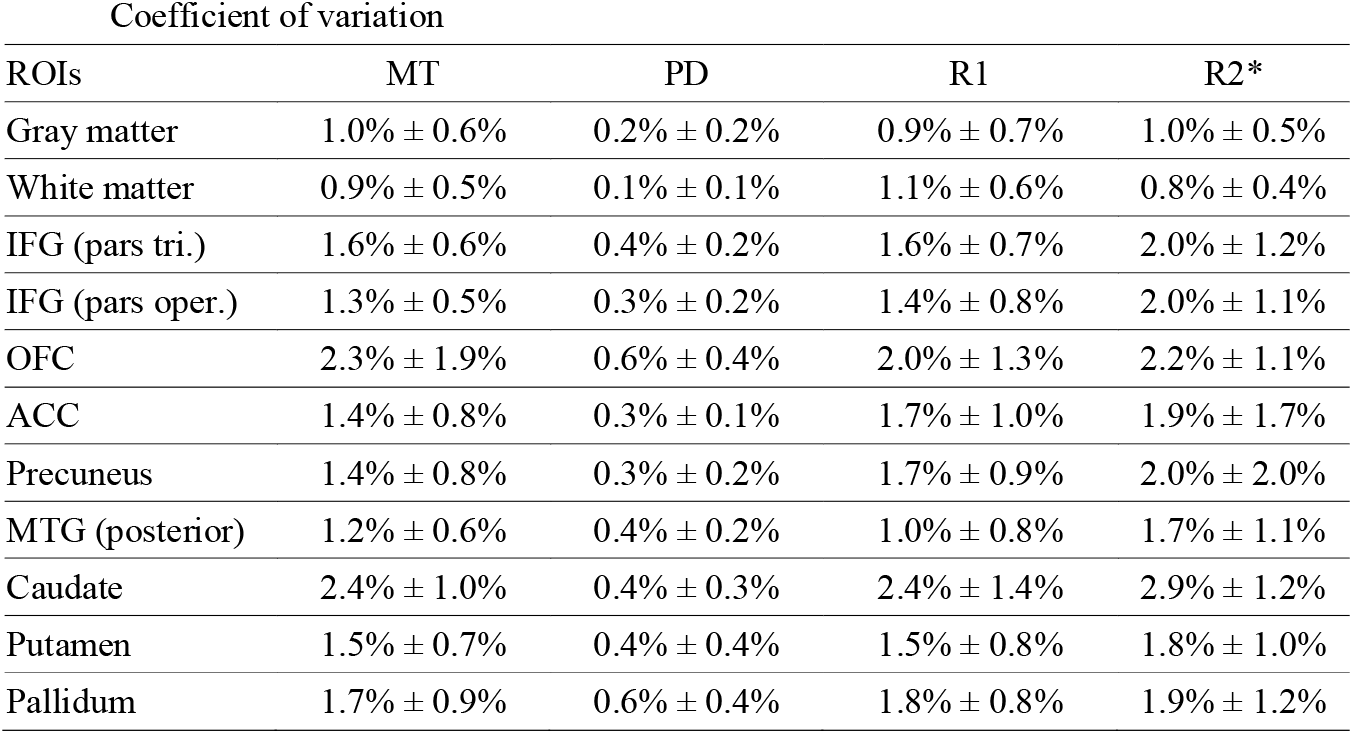
Parameter-specific means and SDs of CoVs across participants, where each participant’s CoV is calculated by dividing the SD across the four measurement points (normalized by *n*−1=3) by the mean of the four extracted values.

| Coefficient of variation |  |  |  |  |
| --- | --- | --- | --- | --- |
| ROIs | MT | PD | R1 | R2* |
| Gray matter | 1.0% ± 0.6% | 0.2% ± 0.2% | 0.9% ± 0.7% | 1.0% ± 0.5% |
| White matter | 0.9% ± 0.5% | 0.1% ± 0.1% | 1.1% ± 0.6% | 0.8% ± 0.4% |
| IFG (pars tri.) | 1.6% ± 0.6% | 0.4% ± 0.2% | 1.6% ± 0.7% | 2.0% ± 1.2% |
| IFG (pars oper.) | 1.3% ± 0.5% | 0.3% ± 0.2% | 1.4% ± 0.8% | 2.0% ± 1.1% |
| OFC | 2.3% ± 1.9% | 0.6% ± 0.4% | 2.0% ± 1.3% | 2.2% ± 1.1% |
| ACC | 1.4% ± 0.8% | 0.3% ± 0.1% | 1.7% ± 1.0% | 1.9% ± 1.7% |
| Precuneus | 1.4% ± 0.8% | 0.3% ± 0.2% | 1.7% ± 0.9% | 2.0% ± 2.0% |
| MTG (posterior) | 1.2% ± 0.6% | 0.4% ± 0.2% | 1.0% ± 0.8% | 1.7% ± 1.1% |
| Caudate | 2.4% ± 1.0% | 0.4% ± 0.3% | 2.4% ± 1.4% | 2.9% ± 1.2% |
| Putamen | 1.5% ± 0.7% | 0.4% ± 0.4% | 1.5% ± 0.8% | 1.8% ± 1.0% |
| Pallidum | 1.7% ± 0.9% | 0.6% ± 0.4% | 1.8% ± 0.8% | 1.9% ± 1.2% |

### Reliability of MPM parameters and variances explained

In Table 3 we summarize all estimates for the four sources of variance: true-score variance (i.e., variance attributable to between-person differences), day-specific error variance, session-specific error variance, and residual error variance, as well as ICC and ICC2, calculated using ICED. As the variance components are rescaled such that they add up to one, they can directly be interpreted as relative contributions to the total observed variance.

**Table 3.** Variance estimates and reliability measures.

| ROI | Modality | RMSEA | Var T | Var D | Var S | Var E | ICC | 95% CI | ICED ICC2 | 95% CI |
| --- | --- | --- | --- | --- | --- | --- | --- | --- | --- | --- |
| Gray matter<br>1 016 729 voxels | MT | 0.215 | 0.883* | 0.000 | 0.003 | 0.114* | 0.883 | 0.759–0.951 | 0.967 | 0.930–0.986 |
|  | PD | 0.449 | 0.873* | 0.000 | 0.021 | 0.106* | 0.873 | 0.636–0.963 | 0.958 | 0.855–0.991 |
|  | R1 | 0.357 | 0.728* | 0.000 | 0.000 | 0.272* | 0.728 | 0.348–0.900 | 0.914 | 0.738–0.975 |
|  | R2* | 0.000 | 0.781* | 0.090 | 0.072 | 0.057* | 0.781 | 0.547–0.866 | 0.885 | 0.740–0.939 |
| White matter<br>498 645 voxels | MT | 0.194 | 0.875* | 0.000 | 0.000 | 0.125* | 0.875 | 0.764–0.943 | 0.965 | 0.912–0.982 |
|  | PD | 0.291 | 0.808* | 0.000 | 0.020 | 0.172* | 0.808 | 0.458–0.914 | 0.937 | 0.682–0.979 |
|  | R1 | 0.386 | 0.733* | 0.000 | 0.029 | 0.238* | 0.733 | 0.385–0.854 | 0.906 | 0.737–0.956 |
|  | R2* | 0.125 | 0.883* | 0.062 | 0.000 | 0.055* | 0.883 | 0.624–0.950 | 0.952 | 0.825–0.986 |
| IFG (pars tri.)<br>9 503 voxels | MT | 0.000 | 0.757* | 0.000 | 0.092 | 0.151* | 0.757 | 0.598–0.864 | 0.891 | 0.769–0.948 |
|  | PD | 0.522 | 0.769* | 0.000 | 0.000 | 0.231* | 0.769 | 0.465–0.903 | 0.930 | 0.807–0.973 |
|  | R1 | 0.274 | 0.562* | 0.000 | 0.000 | 0.438* | 0.562 | 0.242–0.769 | 0.837 | 0.512–0.929 |
|  | R2* | 0.158 | 0.389* | 0.000 | 0.229 | 0.381* | 0.389 | 0.030–0.671 | 0.627 | 0.082–0.902 |
| IFG (pars oper.)<br>11 674 voxels | MT | 0.000 | 0.835* | 0.000 | 0.069 | 0.096* | 0.835 | 0.654–0.914 | 0.927 | 0.816–0.971 |
|  | PD | 0.420 | 0.816* | 0.000 | 0.000 | 0.184* | 0.816 | 0.536–0.926 | 0.946 | 0.783–0.981 |
|  | R1 | 0.122 | 0.547* | 0.000 | 0.000 | 0.453* | 0.547 | 0.173–0.797 | 0.828 | 0.510–0.935 |
|  | R2* | 0.113 | 0.427* | 0.199 | 0.000 | 0.374* | 0.427 | 0.000–0.742 | 0.688 | 0.000–0.902 |
| OFC<br>25 157 voxels | MT | 0.258 | 0.570* | 0.000 | 0.000 | 0.430* | 0.570 | 0.246–0.819 | 0.841 | 0.557–0.953 |
|  | PD | 0.307 | 0.584* | 0.000 | 0.000 | 0.416* | 0.584 | 0.354–0.685 | 0.849 | 0.745–0.900 |
|  | R1 | 0.132 | 0.424* | 0.000 | 0.000 | 0.575* | 0.424 | 0.000–0.778 | 0.747 | 0.042–0.931 |
|  | R2* | 0.290 | 0.523* | 0.000 | 0.134 | 0.343* | 0.523 | 0.202–0.770 | 0.761 | 0.433–0.917 |
| ACC<br>20 844 voxels | MT | 0.192 | 0.817* | 0.000 | 0.000 | 0.182* | 0.817 | 0.630–0.928 | 0.947 | 0.869–0.980 |
|  | PD | 0.541 | 0.828* | 0.000 | 0.000 | 0.172* | 0.828 | 0.693–0.911 | 0.950 | 0.877–0.974 |
|  | R1 | 0.153 | 0.453* | 0.000 | 0.000 | 0.547* | 0.453 | 0.124–0.727 | 0.768 | 0.380–0.917 |
|  | R2* | 0.338 | 0.754* | 0.000 | 0.000 | 0.246* | 0.754 | 0.410–0.919 | 0.924 | 0.737–0.976 |
| Precuneus<br>44 699 voxels | MT | 0.000 | 0.834* | 0.000 | 0.031 | 0.135* | 0.834 | 0.629–0.942 | 0.942 | 0.809–0.978 |
|  | PD | 0.241 | 0.840* | 0.016 | 0.000 | 0.145* | 0.840 | 0.660–0.945 | 0.950 | 0.894–0.983 |
|  | R1 | 0.208 | 0.612* | 0.000 | 0.000 | 0.388* | 0.612 | 0.307–0.803 | 0.863 | 0.623–0.941 |
|  | R2* | 0.304 | 0.209 | 0.626* | 0.000 | 0.165* | 0.209 | 0.000–0.671 | 0.371 | 0.000–0.856 |
| MTG (post.)<br>21 879 voxels | MT | 0.300 | 0.780* | 0.000 | 0.000 | 0.220* | 0.780 | 0.542–0.898 | 0.934 | 0.831–0.973 |
|  | PD | 0.177 | 0.836* | 0.017 | 0.031 | 0.117* | 0.836 | 0.701–0.919 | 0.938 | 0.834–0.976 |
|  | R1 | 0.000 | 0.660* | 0.105 | 0.129 | 0.106* | 0.660 | 0.151–0.875 | 0.810 | 0.378–0.960 |
|  | R2* | 0.173 | 0.532* | 0.297* | 0.000 | 0.171* | 0.532 | 0.234–0.759 | 0.735 | 0.428–0.920 |
| Caudate<br>5 750 voxels | MT | 0.000 | 0.346* | 0.000 | 0.000 | 0.654* | 0.346 | 0.083–0.539 | 0.679 | 0.197–0.822 |
|  | PD | 0.000 | 0.662* | 0.000 | 0.000 | 0.337* | 0.662 | 0.214–0.858 | 0.887 | 0.635–0.967 |
|  | R1 | 0.294 | 0.433* | 0.000 | 0.000 | 0.567* | 0.433 | 0.222–0.656 | 0.753 | 0.594–0.897 |
|  | R2* | 0.000 | 0.571* | 0.000 | 0.000 | 0.429* | 0.571 | 0.183–0.845 | 0.842 | 0.458–0.951 |
| Putamen<br>9 626 voxels | MT | 0.236 | 0.780* | 0.000 | 0.000 | 0.220* | 0.780 | 0.550–0.893 | 0.934 | 0.821–0.969 |
|  | PD | 0.418 | 0.596* | 0.000 | 0.211 | 0.193* | 0.596 | 0.165–0.861 | 0.771 | 0.264–0.957 |
|  | R1 | 0.220 | 0.666* | 0.000 | 0.034 | 0.301* | 0.666 | 0.328–0.851 | 0.876 | 0.618–0.963 |
|  | R2* | 0.000 | 0.766* | 0.077 | 0.014 | 0.143* | 0.766 | 0.227–0.918 | 0.903 | 0.429–0.978 |
| Pallidum<br>4 244 voxels | MT | 0.244 | 0.817* | 0.000 | 0.017 | 0.166* | 0.817 | 0.641–0.917 | 0.941 | 0.857–0.976 |
|  | PD | 0.034 | 0.483* | 0.000 | 0.197 | 0.320* | 0.483 | 0.192–0.741 | 0.710 | 0.339–0.900 |
|  | R1 | 0.213 | 0.715* | 0.000 | 0.000 | 0.285* | 0.715 | 0.338–0.850 | 0.909 | 0.726–0.959 |
|  | R2* | 0.000 | 0.861* | 0.033 | 0.023 | 0.082* | 0.861 | 0.669–0.945 | 0.945 | 0.833–0.985 |

\* = $p$ of chisq difference from null model < .05 (VarT, VarD, VarS), or $p$ of Wald test < .05 (Var E). RMSEA = Root Mean Squared Error of Approximation

#### Variance in MT

In both whole gray and white matter, most variance in MT could be attributed to true score variance (ca. 88%). There were no day- or session-specific effects, and relatively small amounts of variance appeared as residual error (11% and 13%). This was also reflected in high ICC values (GM: 0.88, WM: 0.88). In the localized ROIs, true-score variance in MT varied quite a bit, from high (84% in pars opercularis, 83% in precuneus, 82% in ACC, 82% in pallidum, 78% in MTG, 78% in putamen, 76% in pars triangularis), to lower (57% in OFC), and very low (35% in caudate). There were no significant day- or session-specific effects in any of the regions. Therefore, ICCs of MT were excellent for most of the regions and were only lower for OFC and caudate. In particular, caudate showed a poor reliability estimation of MT compared to the other regions. This is also reflected in the wide confidence interval of the ICC for MT in caudate, ranging from 0.07 to 0.54.

#### Variance in PD

In both whole GM and WM, the relative proportion of true-score variance for PD was very high (87% and 81%), relatively small amounts appeared as residual error variance (11% and 17%) and again no significant effects of day or session appeared. This was also reflected in high ICC values. Note, though, that per construction the absolute variance of PD in WM is negligible as PD is set to 69 p.u., see also second row from Figure 7. True-score variance for PD was also high in precuneus (84%), MTG (84%), ACC (83%), pars opercularis (82%), and pars triangularis (77%), and slightly lower in caudate (66%), putamen (60%), OFC (58%), and pallidum (48%), again with no significant day- or session-specific variances. ICC values for PD were all excellent, except for lower values in caudate (0.66, CI ranging from 0.15 to 0.85), in putamen (0.60, CI ranging from 0.17 to 0.86), in OFC (0.58, CI ranging from 0.41 to 0.70), and pallidum (0.48, CI ranging from 0.19 to 0.74).

**Figure 7.**
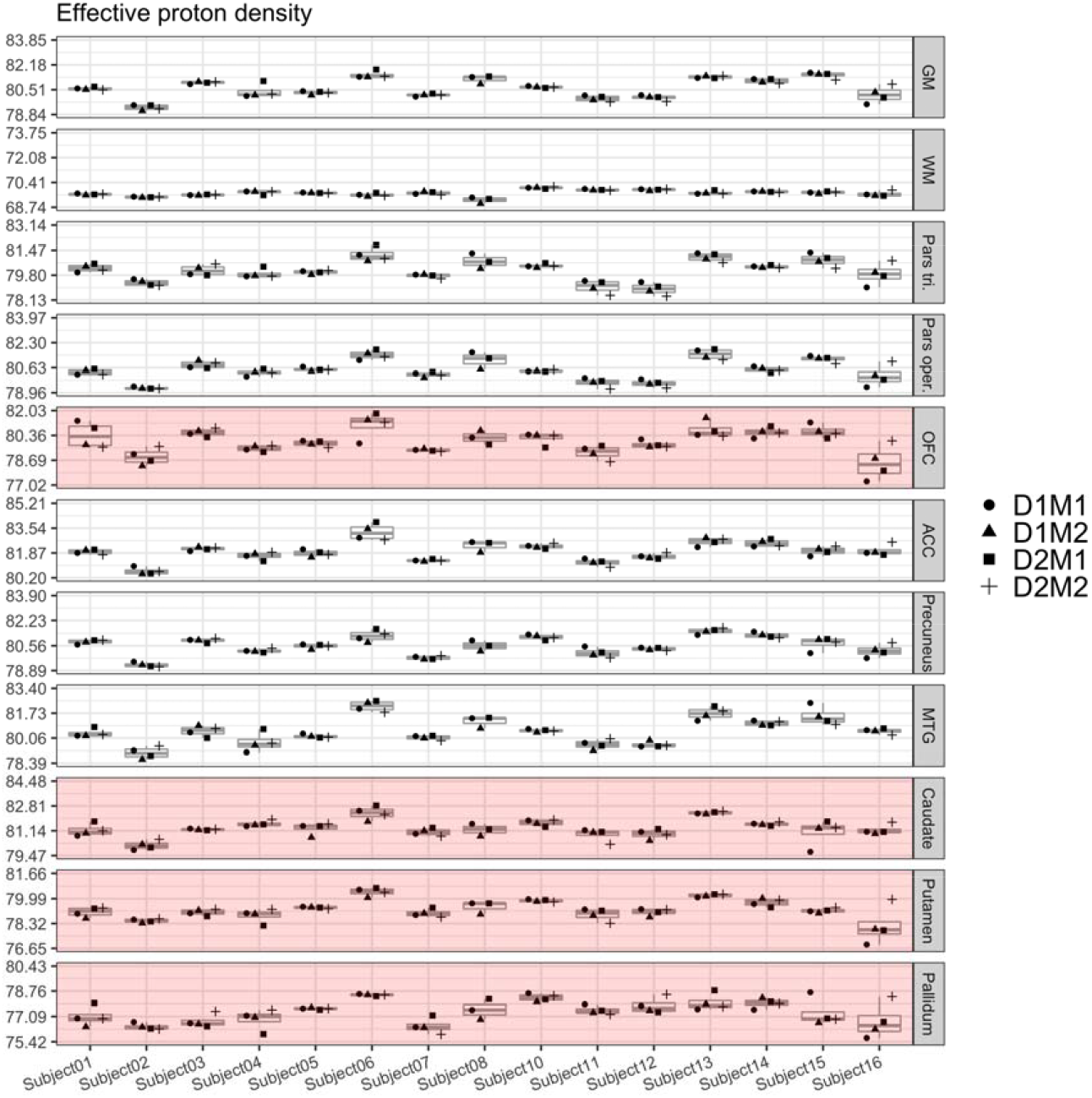
Boxplots of PD values for each of the 15 participants and their four measurements. Parts of the plots marked in red indicate an attenuated ICC value (<0.75) for this region. Note that the range displayed on the y-axis differs across ROIs; importantly, though, the width of the displayed range is constant across ROIs to ensure comparability and is always 5.01.

#### Variance in R1

In general, R1 and R2* values exhibited overall much smaller proportions of true-score variances than MT and PD parameters. In both whole GM and WM, true-score variance for R1 accounted for 73% of the estimated variance, and larger amounts appeared as residual error variance (27% and 24%), while effects of day and session were still not significantly different from zero. Accordingly, ICC values for R1 in whole GM and WM were only fair, both at 0.73. For the smaller, localized ROIs, true-score variances for R1 were considerably smaller, the highest being in pallidum (72%), putamen (67%), MTG (66%), precuneus (61%), pars triangularis (56%), and pars opercularis (55%), and the lower ones in ACC (45%), caudate (43%), and OFC (42%), the latter three with a higher proportion of residual error variance than true-score variance at 55%, 57%, and 58%. Accordingly, ICC values were rather low for ACC, OFC, and caudate with wide confidence intervals and only slightly better for pars triangularis, pars opercularis, precuneus, and MTG. Indeed, for R1, none of the ICC values was above our desired threshold for reliability of 0.75.

#### Variance in R2*

In GM and WM, true-score variance was found to be 78% and 88% of the total variance, with a residual error variance of 6% in each. ICC values were excellent at 0.78 and 0.88. The proportion of true-score variance for R2* was excellent in ROIs of the basal ganglia, namely 77% in putamen, 86% in pallidum, and was also excellent in ACC (75%). However, it was considerably lower in the remaining ROIs, with 57% in caudate, 53% in MTG, 52% in OFC, and 43% in pars opercularis, and was poor in pars triangularis with 39% and in precuneus with 21%, a value not significantly different from the null model. For R2*, there were significant day-specific effects in MTG (Var D = 30%) and precuneus (Var D = 63%). For R2*, ICC values were overall relatively low, compared to MT and PD, except for putamen and pallidum and ACC, and reached an unacceptable value of 0.21 in precuneus.

#### Voxel-specific analysis

In addition to the ROI analysis, we performed the above-described reliability estimation also in every single voxel. Figure 12 displays true score variance, that is identical to ICC in our case, such that the whiter a voxel appears, the higher its ICC value. A visual inspection of these images mirrors the above described pattern of overall good ICC for MT and PD, but attenuated values for R1 and R2*.

## Discussion

In this study, we investigated the test-retest reliability of four MPM parameters, namely MT, PD, R1, and R2*, in whole gray and white matter as well as in selected gray matter ROIs that are commonly of interest in cognitive neuroscience studies. To evaluate reliability, we used ICED (Brandmaier, Wenger, et al., 2018), which partitions multiple sources of unreliability into its constituent components and therefore provides a more detailed picture of the parameter properties than CoV or ICC alone.

A basic check of value plausibility shows that the measured parameter values fall well within the range of those from previously published studies. For example, the R1 = 0.62/0.98 s^-1^ in GM/WM was similar to the R1 reported previously 0.61/1.04 s-1 (Weiskopf et al., 2013) and 0.63/1.19 s^-1^ (Wright et al., 2008). The same holds for our PD estimates of 80.6/81.3 p.u. in GM/caudate, which were comparable to previous studies reporting 84.4/82.7 p.u. (Weiskopf et al., 2013), 81.1/81.5 p.u. (Volz et al., 2012), and 82.2/84.8 p.u. (Neeb, Ermer, Stocker, & Shah, 2008) for the same structures. The small numerical difference here might originate from the usage of effective PD, that is, not extrapolating to TE = 0 in some previous publications as for example in Weiskopf and colleagues (2013). Our estimates of R2* were also similar to previously published values, namely 16.9/21.5 s^-1^ compared to 15.2/21.0 s^-1^ in GM/WM (Weiskopf et al., 2013) and 19.5 s^-1^ and 21.7 s^-1^ in WM (Baudrexel et al., 2009; Martin, Wieler, & Gee, 2008). For all parameters, the choice of resolution can lead to a certain change of the measured parameter values by partial volume effects. In case of R2*, the chosen resolution can produce even more substantial deviations due to the fact that for larger voxels the intra-voxel B_0_ field is more inhomogeneous, which causes larger R2* values.

All four MPM parameters showed excellent reliabilities for whole gray and white matter across the four measurements. However, we noted marked differences in reliability among the four MPM parameters for different regions of the brain: MT and PD exhibited excellent reliability and were robust against participant repositioning within a scanning session and on different days in nearly all regions, except for OFC (ICCs fair at 0.57 for MT and 0.58 for PD), caudate (ICC for MT poor at 0.35, ICC for PD good at 0.66), pallidum (ICC for PD fair at 0.48), and putamen (ICC for PD good at 0.60). In contrast, R1 and R2* showed only fair reliability in most regions (< 0.60) with the exception of regions of the basal ganglia, namely putamen and pallidum, and even poor reliability (< 0.40) for R2* in pars triangularis and precuneus. In some regions, the sum of the error estimates exceeded between-person differences (e.g., for R1 in ACC, OFC, and caudate), effectively rendering it hard if not impossible to interpret between-person differences and continue with correlational approaches. For R2* in precuneus, the proportion of true-score variance was not significantly different from zero, that is, all participants yielded very similar values on this parameter in precuneus. For that region, R2* did not convey any person-specific information that would be potentially associated to any sort of between-person differences in behavior. Given that cognitive neuroscience often aims at delineating brain-behavior relations (Blakemore & Lindenberger, 2020; Krakauer, Ghazanfar, Gomez-Marin, MacIver, & Poeppel, 2017), knowing about these differences in reliability is of critical importance.

These observations on differential reliability of the four MPM parameters reported above exist in parallel to observations of high precision of these parameters, when calculated as the variation over four different measurements on the same scanner, with CoVs ranging between 0.2% and 2.4%. Our estimates of CoVs are even better than those reported before, for example, in an inter- and intra-site validation study (Weiskopf et al., 2013), or in another study testing within-site and between-site reproducibility of MPM (Leutritz et al., 2020). In general, other studies have also found quantitative MRI to be highly reproducible across software updates, different sites, and even across different vendors (Gracien et al., 2020; Lee, Callaghan, Acosta-Cabronero, Lutti, & Nagy, 2019; Leutritz et al., 2020). It is important to note that the findings of high reproducibility as assessed with CoVs, and those of varying reliability between the four MPM parameters across different ROIs as assessed with ICCs do not contradict each other, but are simply different pieces of the same puzzle. To repeat, these two notions of reliability both capture the precision of measurement, however they standardize them with respect to different sample values; CoV standardizes against the sample mean, whereas ICC standardizes against the between-person variance. In experimental approaches, the former logic may be more apt to quantify a standardized precision of measurement, whereas the precision necessary for correlational (individual differences) studies is better captured by the latter.

By definition, ICC values must be interpreted contingent upon the characteristics of a given population. This is important to keep in mind when interpreting a given ICC. To the extent a given study population has lower or higher true score variability, reliability will increase of decrease proportionally. Here, we have measured 15 healthy younger adults aged 20 to 30 years. In this sample, ICCs were high for MT and PD but attenuated for R1 and R2*. It is an empirical question whether similar results would be obtained in samples representing an older or diseased population. For example, it may very well be that the underlying biological characteristics associated with these parameters simply do not yet show or do not anymore show any substantial variance in the 15 younger adults included in this study as childhood and adolescence is over and older age has not yet begun. Also, the chosen ROI approach here might limit the ability of MPM to pick up small changes. Classical voxel-based mass univariate mapping approaches might be better suited to deliver more fine-grained results that can be picked up at the individual voxel level, but go unnoticed in the averaging process done to form an ROI.

According to a commonly accepted biophysical interpretation, R1 depends on the mobility of water in its microenvironment, which is affected by certain aspects of cell membranes, and is thus related to myelination due to the presence of dense multiple myelin sheaths (Weiskopf et al., 2021). R2* is dependent on the magnetic field distribution, which in turn is particularly affected by iron (Cherubini, Péran, Caltagirone, Sabatini, & Spalletta, 2009; Duyn et al., 2007; Péran et al., 2009, 2007; Weiskopf et al., 2021). Healthy young adults may simply not differ substantially in iron deposition and molecular mobility near membranes. In other words, there may not have been between-person differences to be detected by R1 and R2* in this specific sample. The fact that reliability of R2* was considerably higher in regions of the basal ganglia speaks to the fact that this part of the brain is indeed specifically amenable to iron deposition and may therefore also be more likely to show individual differences in R2* than other cortical regions. Future studies using ICC as a reliability measure of MPM parameters in more diverse samples will show whether reliability attenuation of R1 and R2* in most cortical regions generalizes beyond the present sample of healthy young adults.

The importance of considerations on reliability of different parameters cannot be overstated. As statistical power is directly influenced by precision of measurement (besides sample size, test size, and population effect size), it is essential to be aware of the reliability of each parameter that is used to characterize, for example, age-related changes in brain structure, as is often intended when using the MPM protocol. With knowledge of reliability estimates, it is then possible to perform power analyses in the context of individual differences in longitudinal designs (Brandmaier, von Oertzen, Ghisletta, Lindenberger, & Hertzog, 2018), thereby enabling informed study design decisions to optimize conditions to detect a hypothesized effect. The “replicability revolution” in psychological science is an example of how changing norms can shape research practices and standards. In only a few years, practices to foster replicability, be it preregistration of hypotheses and planned analyses or publishing of analysis scripts or even data, have rapidly gained in popularity (Nosek, Ebersole, DeHaven, & Mellor, 2018). Similar norms would be highly desirable in the context of reliability: Researchers should report the reliabilities of all MRI-derived estimates whenever these are used to study individual differences, similar to what has been proposed for task-fMRI (Elliott et al., 2020). In doing so, researchers can make more informed decisions about sample size and select reliable measures for a given research question.

At the risk of redundancy, we would like to reiterate that all four MPM parameters showed excellent reproducibility, that is, they show very little variability when measurements are repeated in the same participant four times. However, the four parameter estimates do differ in how informative they are for assessing between-person differences in young adults. In particular, R1 and R2* seem to carry only limited person-specific information as can be gathered from the very low variability among participants displayed in Figures 8 and 10.

**Figure 8.**
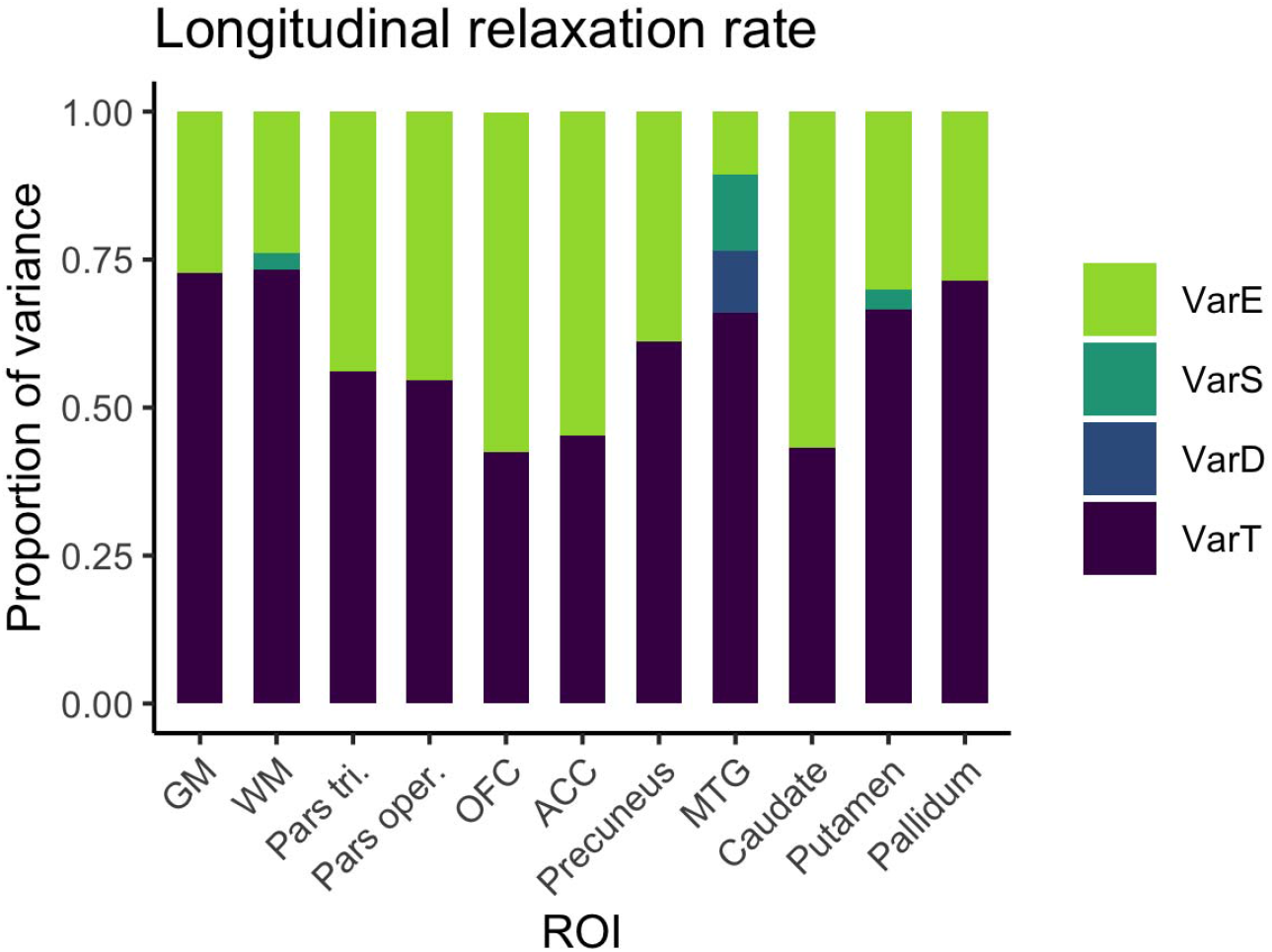
Distribution of magnitudes of sources of variance for R1 across the different ROIs. Note that absolute magnitudes are displayed, irrespective of significance. Also for R1, session- and day-specific variances were not significantly different from zero.

**Figure 9.**
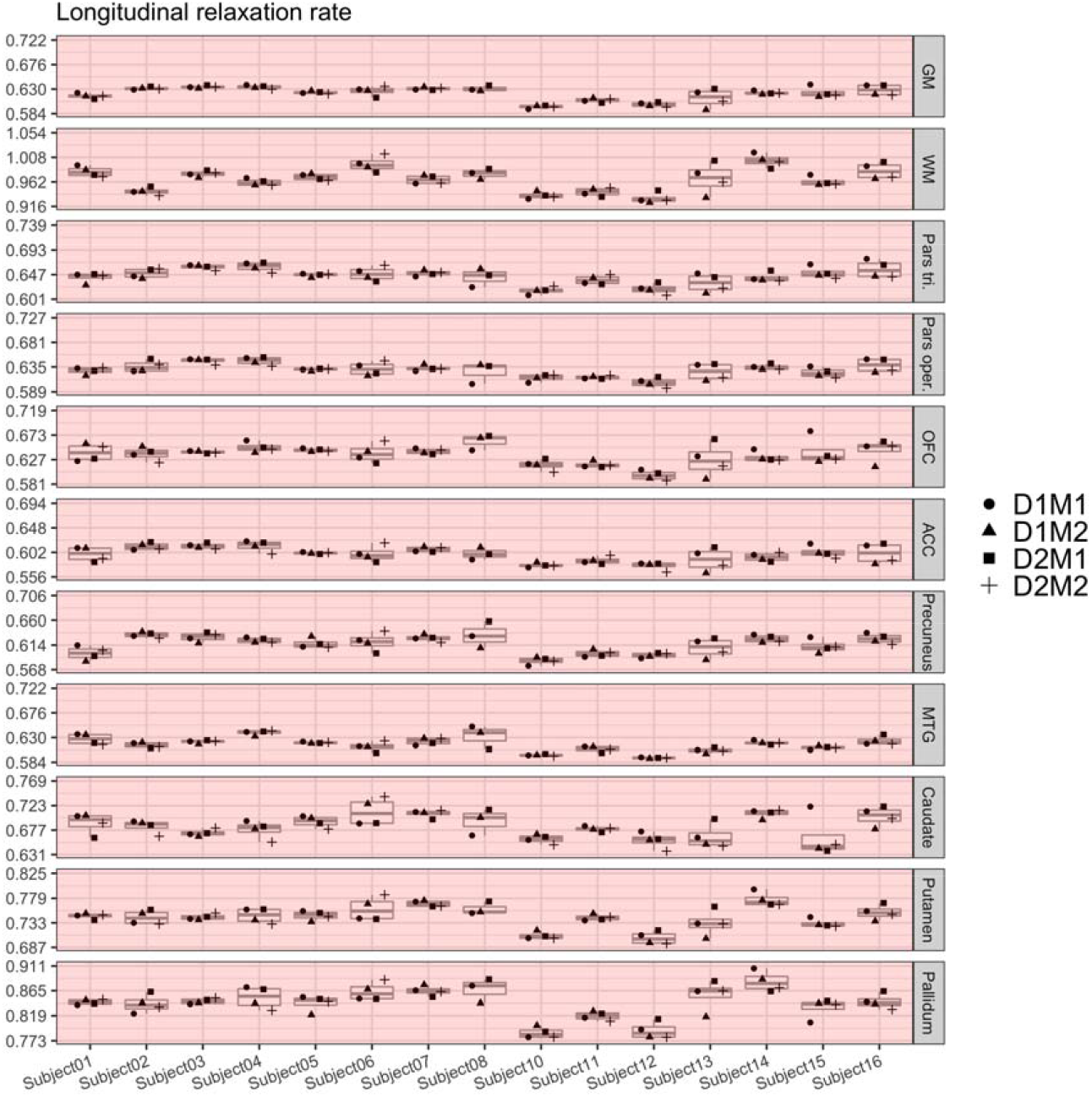
Boxplots of R1 values for each of the 15 participants and their four measurements. ICC values were below 0.75 in all investigated regions (marked in red). Note that the range displayed on the y-axis differs across ROIs; importantly, though, the width of the displayed range is constant across ROIs to ensure comparability and is always 0.138.

**Figure 10.**
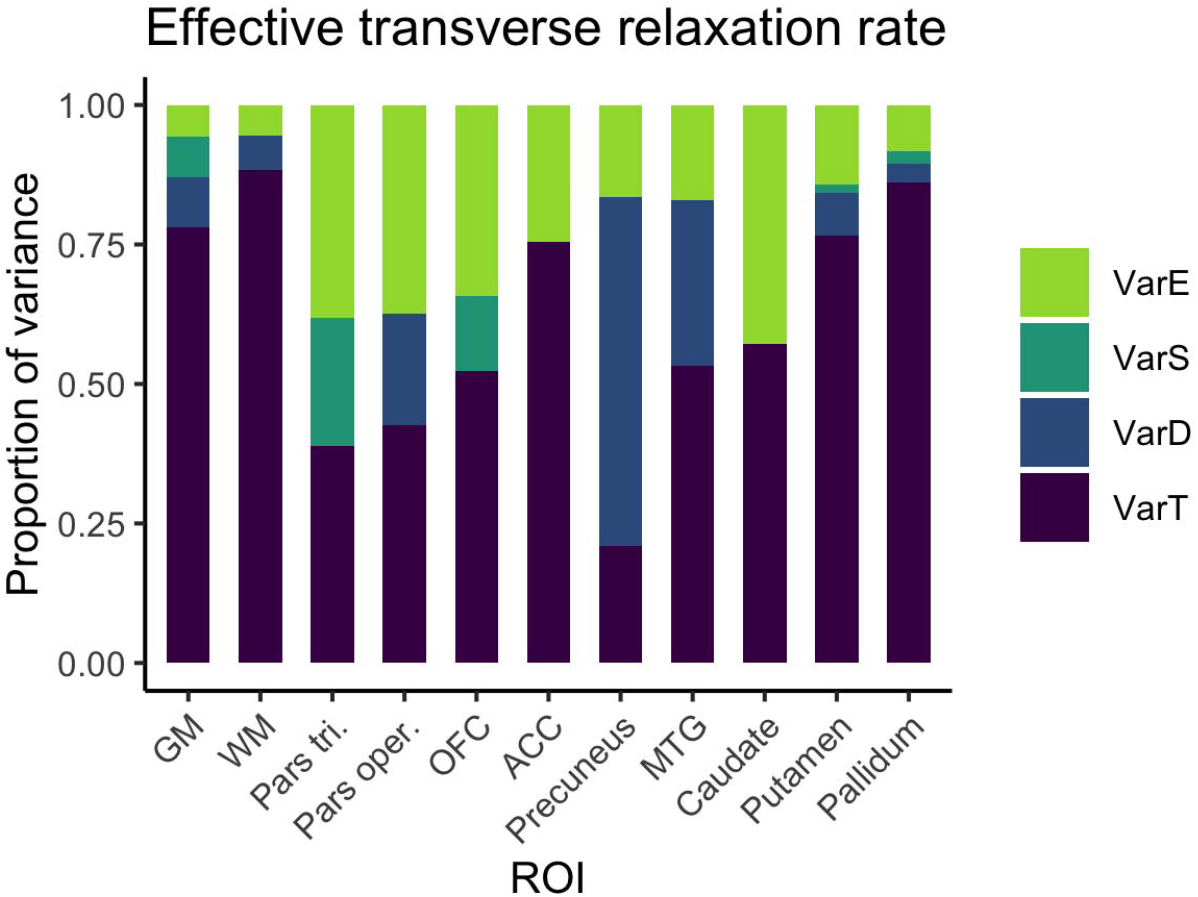
Distribution of magnitudes of sources of variance for R2* across the different ROIs. Note that absolute magnitudes are displayed, irrespective of significance. Notably, there were significant day-specific variances for R2* in MTG and precuneus.

**Figure 11.**
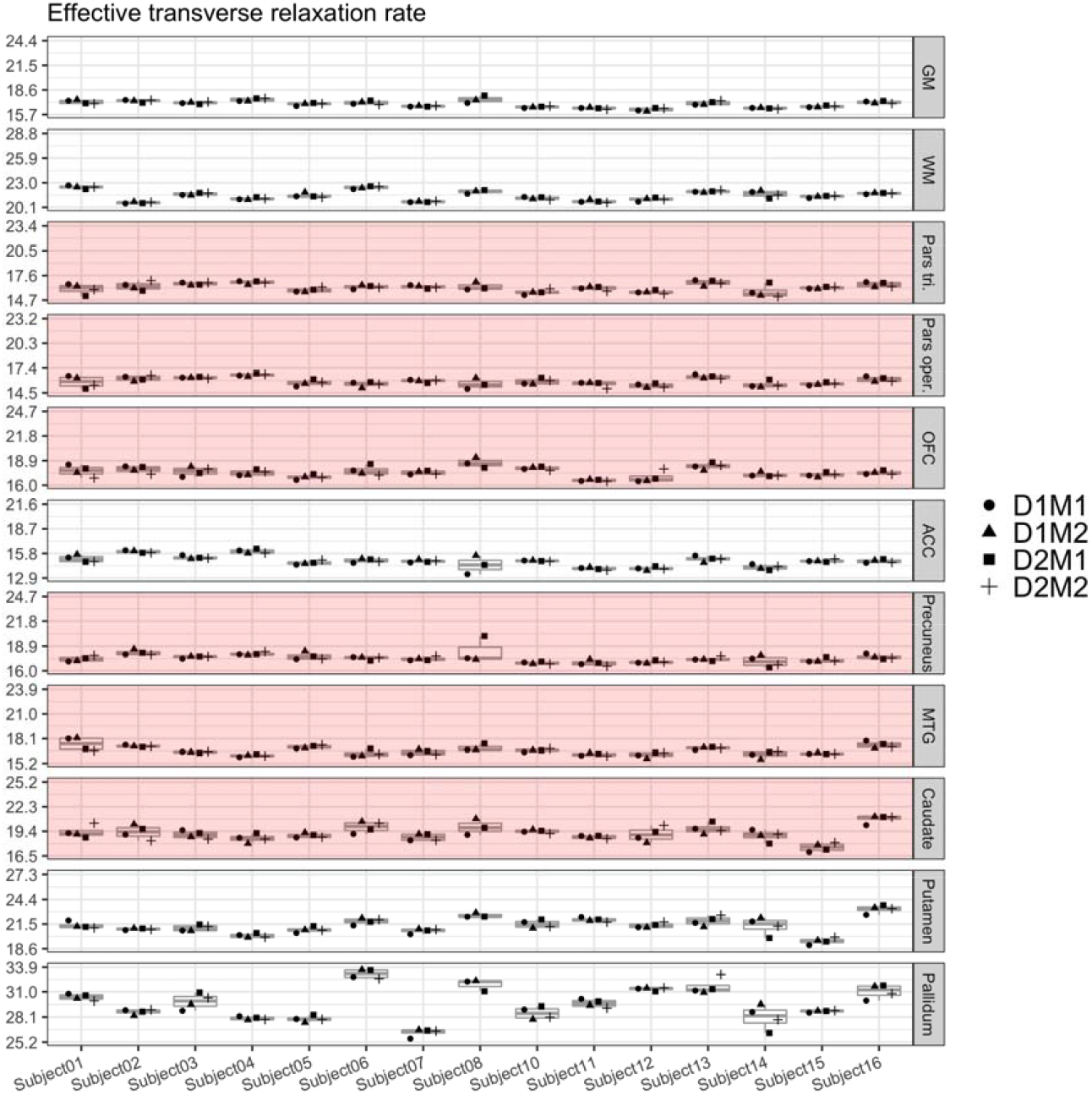
Boxplots of R2* values for each of the 15 participants and their four measurements. Parts of the plots marked in red indicate an attenuated ICC value (<.75) for this region. Note that the range displayed on the y-axis differs across ROIs; importantly, though, the width of the displayed range is constant across ROIs to ensure comparability and is always 8.7.

**Figure 12.**
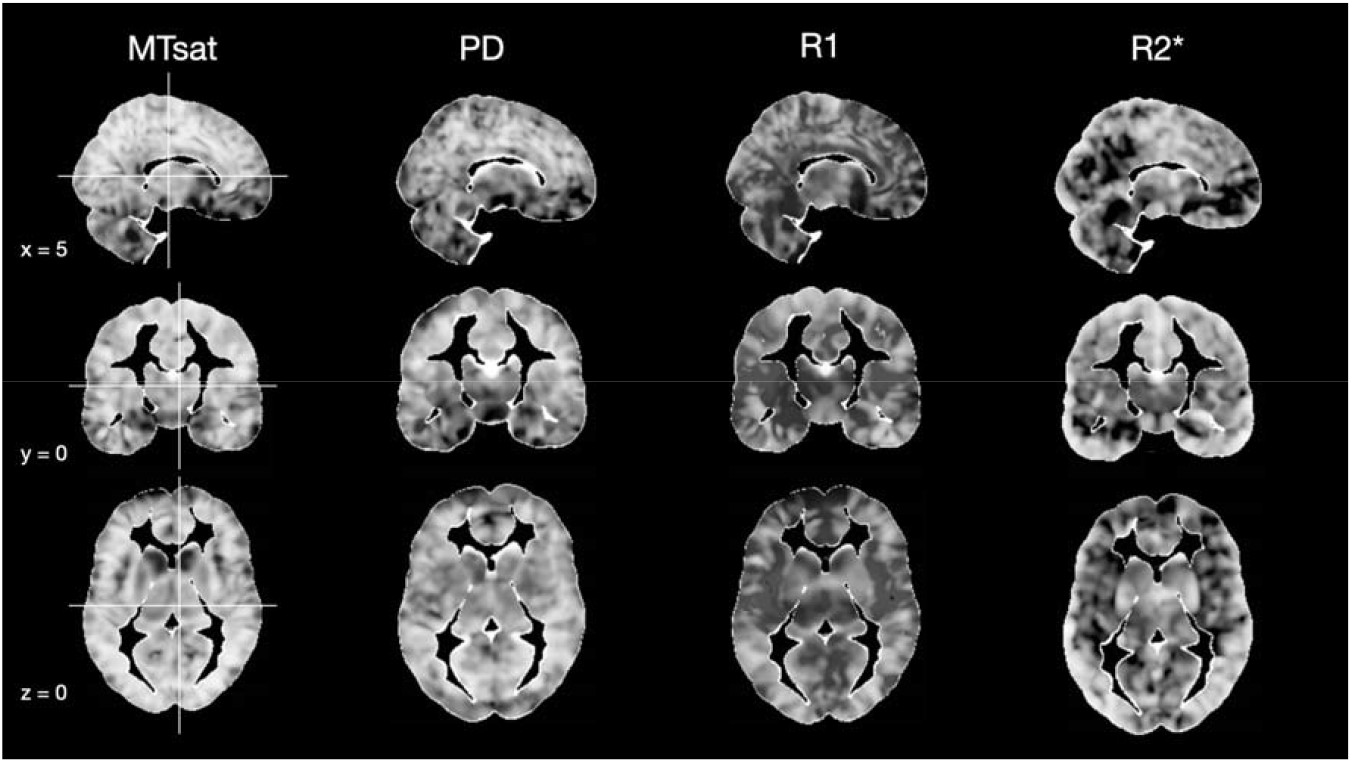
Voxel-specific estimation of true score variance, that is, ICC, for all four MPM parameters. The whiter a voxel appears, the higher its ICC value.

In the following, we discuss this differential reliability in light of a number of potential influences on parameter estimation.

### Potential causes of variations in parameter maps

The dual flip angle mapping approach used in MPM (Helms, Dathe, & Dechent, 2008) provides signal amplitude (proportional to PD) and R1 maps that need to be corrected for RF transmit and receive field inhomogeneities. As recommended for the preprocessing stream, we used highly accurate and precise RF transmit field maps with a total error of less than 3% (Lutti et al., 2010, 2012) and corrected for imperfect RF spoiling. The previous effects lead to deviations from the Ernst signal equation underlying the R1 estimation and imprecisions in their correction may thus lead to inaccuracies (Corbin & Callaghan, 2021; Preibisch & Deichmann, 2009; Yarnykh, 2010). For example, even small deviations of 3% in the RF transmit field mapping can cause errors of up to 6% in the R1 maps due to the quadratic dependence of the estimated R1 on the local flip angle (Weiskopf et al., 2011). MT saturation maps are largely self-correcting and independent of the RF transmit and receive fields (Helms, Dathe, Kallenberg, et al., 2008). In addition, residual effects were further reduced in post-processing based on the measured RF transmit maps (Weiskopf et al., 2013).

There is some bias to be expected in R2* (and PD) maps as the mono-exponential decay model applied by the ESTATICS approach and extrapolation to TE=0 is just a simplification that does not perfectly fit due to partial voluming of different cell compartments with different transversal relaxation rates as intra- and extra-cellular and myelin-associated spaces. Thus R2* is poorly defined and also the extrapolation of the signal to TE=0 for the PD mapping is inaccurate (Neeb et al., 2008; Tabelow et al., 2019; Weiskopf et al., 2013). Thus, differences in shim, which can also be caused by differences in relative orientation of the head, may have influenced R2* maps (Draganski et al., 2011). R2* of highly structured brain structures can also exhibit a field orientation dependence due to their microstructural geometry even independent of shim quality (Papazoglou et al., 2019; Wharton & Bowtell, 2012). At the same time, the high spatial resolution of 1 mm should reduce the effects of susceptibility artifacts on the signal decay due to a smaller within voxel spin phase coherence loss (Weiskopf, Hutton, Josephs, Turner, & Deichmann, 2007). Also, non-linear B_0_ inhomogeneities may have an influence on the R2* maps, as 3D R2*-mapping by short TE trains have been shown to be affected by local B_0_ gradients (Helms & Dechent, 2010). Since the longest echo time acquired in the PD-weighted sequence was 18.79 ms, the estimation of the long T2* (=1/R2*) found in GM, WM, or CSF is relatively poorly conditioned. The precision of the R2* maps may be improved by increasing the maximal echo time, but this would also prolong the total acquisition time (Weiskopf et al., 2013). Additionally, the generally relatively poor reproducibility and performance of shimming routines might influence the reliable estimation of R2* (Leutritz et al., 2020).

The RF receive field effect on the PD map was minimized by image post-processing. Unified segmentation (Ashburner & Friston, 2005) was adapted to robustly determine and correct for the multiplicative receive coil sensitivity profile in the PD maps, similar to the previously developed UNICORT approach for correcting R1 maps (Weiskopf et al., 2011). Indeed, this seems to be working quite well, as the PD estimates exhibited consistently high ICC values across all the regions in our study (with only a slight attenuation in OFC and caudate).

As with any other MR sequence, MPM performance may be impaired in non-compliant volunteers. For example, some participants may have difficulties to minimize head or body motion, which can change the magnetic field in the head and affect data quality (Versluis et al., 2010; Weiskopf et al., 2013). The parameter maps are estimated from three acquired FLASH sequences and are sensitive to existing artifacts in any of these. When inspecting our plots above depicting the individual means of all four measurements for every participant, it is obvious that some participants’ data was in general more variable than others (e.g., participants 8 and 13). Some of these problems may be alleviated by using prospective motion correction (Callaghan et al., 2015; Maclaren et al., 2012) and phase navigator techniques (Versluis et al., 2010).

We also note that the physical and biophysical models underlying the modeling, analysis and interpretation of quantitative MRI and MPM data pose additional constraints. The brain tissue is a highly complex structure with a plethora of cells, cellular process and extensive vascularization. Thus, e.g. the reduction and description by single compartments underlying standard relaxation parameters such as R1 and R2* can only partially capture the tissue’s complexity, effectively causing instabilities in the aggregate parameter measures (Weiskopf et al., 2021).

### Conclusion

This study used ICED (Brandmaier, Wenger, et al., 2018) to investigate the reliability of multiparameter mapping (MPM) parameters assessed with 3T MRI. ICED allowed us to separate sources of unreliability due to session- and day-specific effects from residual error variance. In line with earlier validation studies, we found high reproducibility of all four MPM parameters using CoV throughout all assessed regions of the brain. Going beyond common practice, we placed special emphasis on representing the precision with which MPM parameters capture between-person differences. To this end, we calculated the ICC based on ICED, which quantifies variance within persons in relation to total variance. We found that reliabilities of between-person differences were high for all four parameters in relation to whole gray and white matter. However, across different regions of the brain, the reliabilities of the four parameters varied greatly. Specifically, MT and PD emerged as highly reliable parameters that are robust against participant repositioning in nearly all regions, whereas true-score variances were lower for R1 and R2*. In some regions, residual-error variances of R1 and R2* exceeded true-score variances, rendering the interpretation of between-person differences for these parameters unviable. We conclude that R1 and R2* carried little personspecific information in regions outside the basal ganglia in the present sample of healthy young adults, and recommend researchers to routinely check the reliability of MRI parameters before examining their associations to individual differences in behavior.

## Acknowledgments

This work was supported by the Max Planck Society and the Max Planck Institute for Human Development and is part of the BMBF funded EnergI Consortium (01GQ1421B). NW was supported by: the European Research Council under the European Union’s Seventh Framework Programme (FP7/2007-2013) / ERC grant agreement n° 616905; the European Union’s Horizon 2020 research and innovation programme under the grant agreement No 681094; the BMBF (01EW1711A & B) in the framework of ERA-NET NEURON. The Wellcome Centre for Human Neuroimaging is supported by core funding from the Wellcome [203147/Z/16/Z]. We are very grateful to Siawoosh Mohammadi for advice on data preprocessing, to Michael Krause for continuous assistance in implementation of data preprocessing on the computing cluster, and to Martina Callaghan for continued support during implementation of the MPM protocol and highly valuable comments on the final manuscript draft. We thank the MRI team at the Max Planck Institute for Human Development (Sonali Beckmann, Nadine Taube; Thomas Feg, and Davide Santoro) as well as all participants for their time and support.

## Data Availability Statement

The data that support the findings of this study as well as the R analysis script are openly available in OSF at https://osf.io/6p9bf/.

